# Simultaneous EEG-fMRI investigation of sound sequence processing in human neonates

**DOI:** 10.64898/2026.01.09.698718

**Authors:** Parvaneh Adibpour, Vyacheslav Karolis, Ines Tomazinho, Dario Gallo, Cidalia Dacosta, Kamilah St Clair, Wendy Norman, Kathleen Colford, Fraser Aitken, Claire Kabdebon, Jonathan O’Muircheartaigh, Tomoki Arichi

## Abstract

The ability to detect sensory regularities and their violations (i.e. deviance) is fundamental for learning and the orientation of attention across the lifespan. This starts from the prenatal period, where the environment is rich in regular patterns and deviations of auditory stimulation which likely provides the basis for the emergence of early learning capacities. Following birth, EEG recordings of brain activity have demonstrated that neonates detect violations of regularity and show a robust mismatch response to deviant auditory stimuli. However, it is unknown how this processing is integrated across wider regions within emergent brain networks.

To gain comprehensive insight into both the temporal and spatial properties of the processing underlying sensory deviance detection in the neonatal brain, we acquired simultaneous EEG and fMRI data in sleeping neonates during an auditory oddball paradigm. A mismatch response was identified with EEG when a deviant sound appeared after a sequence of identical sounds but not when a sound was omitted, suggesting encoding of deviances that violated local sequence regularity, but not global sequence structure. Simultaneous fMRI demonstrated engagement of distributed cortical networks during sound sequence processing, encompassing temporal, sensori-motor, and inferior frontal cortices, as well as subcortical regions including the thalamus and hippocampus. Deviance detection was associated with activity in the superior and ventromedial (hippocampus and parahippocampus) parts of the temporal lobe, and the precentral gyrus. Together, these findings suggest that neonatal brain detects deviance in sound sequences through the encoding of local regularities, supported by distinct processing pathways within sensory and limbic systems. These insights help advance our understanding of early auditory learning mechanisms, with implications for understanding how atypical developmental trajectories may emerge in these substrates.

## Introduction

Infancy is a period of remarkable experiential discovery, during which the developing brain must quickly adapt to make sense of continuous streams of environmental sensory input. The prenatal auditory environment provides a rich tapestry of regular patterns and instances of deviations— embedded in the maternal heartbeat or speech as they travel through the maternal tissue. This early milieu likely helps to prepare early brain networks for detecting regularities and processing violations of regularities (Gervain, 2018; Ghio et al., 2021), a capacity that is fundamental for learning and for orienting attention across the lifespan (Sara & Bouret., 2012; Soltani & Izquierdo, 2019). In doing so, it lays the foundations for later emergence of cognitive abilities such as language learning (Goswami, 2022).

Shortly after birth, neonates are already capable of demonstrating sensitivity to auditory regularities and appearance of deviance. Behavioral reports indicate that newborns can perceive and discriminate rhythm – a form of regularity- in language categories (Nazzi et al., 1998; Ramus et al., 2000). At the neural level, electroencephalography (EEG) and Magnetoencephalography (MEG) studies show that term- and preterm-born neonatal brain activity tracks regularities embedded in auditory streams (Flo et al., 2022; Saadatmehr et al., 2025; Teinonen et al., 2009; Corvilain et al., 2025). Consequently, when these regularities are violated, they show characteristic mismatch responses that are distinct from adults (Dehaene-Lambertz & Pena, 2001; Mahmoudzadeh et al., 2017). Interestingly, neonates are able to detect both simple and complex forms of violations in sound sequence regularity, demonstrated by a mismatch response to novel items within repeated sequences (Mahmoudzadeh et al, 2017) as well as to higher-order deviations (Moser et al.,2020; Panzani et al., 2023) such as disruption to an alternating sequence of sounds. Yet, it is unclear to what extent neonates can encode abstract rules about sequence regularity, and whether they can detect deviations that violate global sequence structure, such as the expected number of elements in a sequence. Addressing this question is crucial for understanding the mechanisms that support early encoding abilities.

Moreover, although EEG and MEG studies indicate the presence and the time-course of responses in neonatal brain, they lack the spatial resolution and sensitivity to activity in the deeper areas of the brain needed to study how developing cortical and subcortical networks are involved in early sound sequence processing and deviance detection. By measuring the Blood Oxygen Level Dependent (BOLD) contrast as an indirect measure of neural activity, functional Magnetic Resonance Imaging (fMRI) offers substantially higher spatial resolution, enabling more precise mapping of whole brain activity. A growing body of work demonstrates the feasibility of studying infant auditory processing with fMRI in the noisy environment of the scanner. These studies have localized functional responses along key auditory and language-related regions to continuous speech (Dehaene-Lambertz et al., 2010; Perani et al., 2011; Shultz et al., 2014), music (Perani et al., 2010; Kosakowski et al., 2023), the human voice (Dehaene-Lambertz et al., 2010; Adam-Darque et al., 2020), and salient and rare noise events (Dall’Orso et al., 2021; Sylvester et al., 2021; Moser et al., 2025). The reported activations span primary and higher-order auditory cortices—including Heschl’s gyrus and the planum temporale, the inferior and middle frontal cortex, in addition to subcortical structures such as the hippocampus, amygdala, as well as the brainstem and cerebellum. Together, these findings highlight early-emerging, functionally pre-specialized networks for processing speech, music, and salient sounds.

Here, we aimed to characterise the precursor mechanisms that enable newborn infants to learn from their early auditory environment by studying the networks supporting structured sound sequence processing and deviance detection. Investigating these processes at a time when brain networks are being established is also relevant for understanding the pathophysiology underlying neurodevelopmental conditions, as emerging evidence suggests that such mechanisms are altered in autism and schizophrenia (Huang et al., 2023; Zhang et al., 2025). To probe the early neural mechanisms that support early deviance detection, we acquired simultaneous EEG-fMRI data in sleeping neonates and examined responses to two types of violations of sound sequence regularity: first-order *local* violations introduced by a change in the immediately repeated sound (as in XXX<u>Y</u>), and higher-order *global* sequence-level violations induced by the omission of an expected sound (as in XXX___). In line with prior neurophysiological evidence, we anticipated that neonates show mismatch response to local violations of regularity. We also hypothesized that if neonates can represent global sequence structure, they would also show a mismatch response to the omission of sounds. By complementing EEG responses with fMRI, we expected to gain a refined spatial characterisation of the underlying functional networks beyond broadly localised activations over the temporal and frontal areas for detection of regularities (Flo et al., 2019; Gervain et al., 2008; Nallet et al., 2023) and local deviations (Mahmoudzadeh et al., 2013; Cabrera & Gervain, 2020) reported in functional Near-Infrared Spectroscopy (fNIRS) studies. If present and in-line with adult fMRI work (Bekinschtein et al., 2009; Al Roumi et al., 2023; Mazancieux et al., 2023), we expected sound-sequence processing and deviance detection to recruit a distributed network encompassing both the primary and non-primary auditory cortices as well as the heteromodal associative regions and subcortical regions.

## Methods

### Procedure

The study was approved by the NHS research ethics committee (REC code: 12/LO/1247) and all data was acquired with written informed parental consent. Prior to scanning, molded dental putty was placed in the external ear (President Putty, Coltene Whaledent, Mahwah, NJ, USA) and adhesive earmuffs (MiniMuffs, Natus Medical Inc., San Carlos, CA, USA) were placed on top to attenuate scanner noise. All neonates were studied during natural sleep, after they were fed and settled.

### Participants

We acquired simultaneous EEG and fMRI data in 25 full-term born neonates. Three neonates’ data were rejected due to heavy motion-corruption. The study group therefore included 22 neonates with a median gestational age at birth of 39.4 weeks (range: 37.3-41.4), and a median postmenstrual age (PMA) at the time of scanning of 39.9 weeks (range: 37.9-44.1); nine were female.

### Paradigm

An adapted auditory oddball paradigm was implemented using blocks of sound sequences presented through MRI compatible headphones (Optoacoustics Ltd, Moshav Mazor, IL), interleaved with periods of rest (i.e. no stimulation) for over 20 minutes. Sound stimulation consisted of two consonant–vowel syllables (/ba/, /kou/), chosen to be distinctive from the background scanner noise. They were presented in sequences of four sounds, in the following patterns: *standard sequence* with four identical sounds (/ba/ /ba/ /ba/ /ba/); *deviant sequence* with a change occurring in the last sound (/ba/ /ba/ /ba/ /kou/); or *omission sequence* with the last sound being omitted (/ba/ /ba/ /ba/ /--/). Within each sequence, there was an interval of 600 ms between the onset of each sound; with an interval of 1500 ms between the fourth sound offset and the onset of the next sequence.

Each block consisted of six sequences with either repetitions of the same sequence (6 standard sequences of the same repeated sound) or violation of regularity in three out of the six sequences (3 deviant or omission sequences interleaved with 3 standard sequences (Figure 1)). A total of 21 blocks were presented, out of which 9 blocks included only standard sequences, 6 included deviant sequences and 6 included omission sequences. The order of presentation of six sequences in deviant/omission blocks was pseudorandomized, always starting with a standard sequence followed by the deviant/omission sequence and randomized for the next four sequences. Each stimulation block lasted 19.92 seconds, followed by at least 24 seconds of rest before the start of the next block. For one infant, omission blocks were replaced by deviant blocks due to technical issues; this infant was therefore excluded from the analyses of omission sequences.

**Figure 1.**
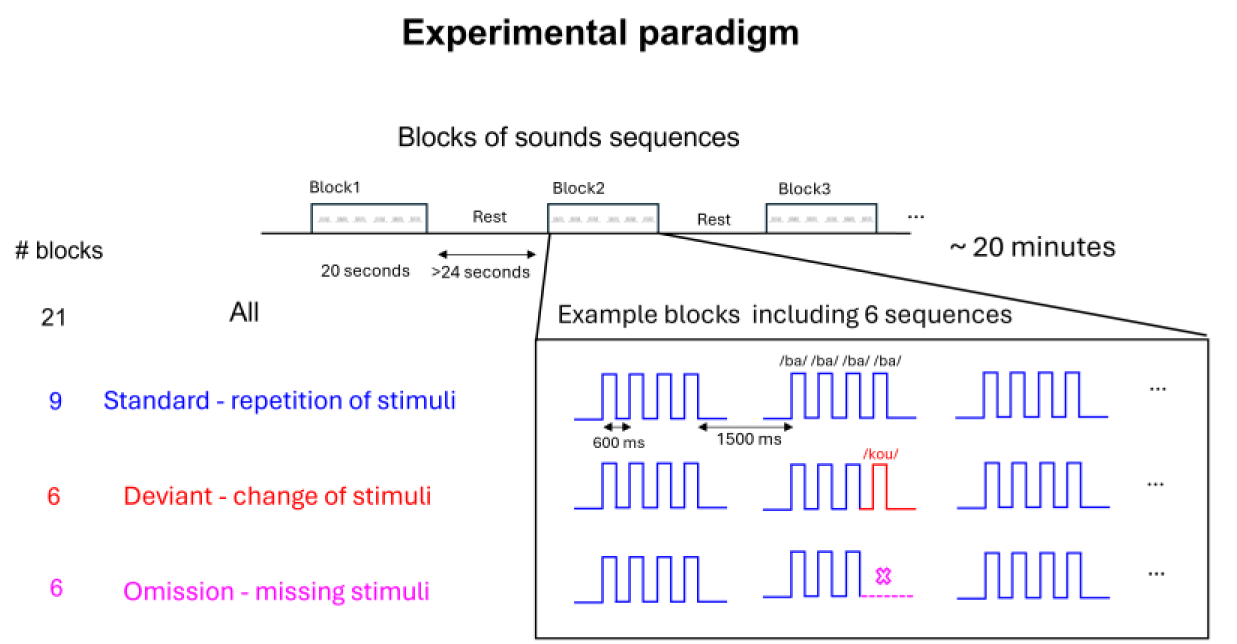
Illustration of the auditory oddball paradigm. Neonates were presented with blocks of sounds interleaved with no-stimulation periods. Each block contained six sound sequences. In standard blocks, all six sequences involved four repetitions of the same sound. In deviant and omission blocks, three of six sequences ended with either a deviant sound or an omitted final sound.

Stimulation presentation was controlled via Matlab and psyhchtoolbox and sound presentations were timestamped on EEG recordings using a hardware solution (TriggerBox, Brain Products, GmbH, Gilching, Germany).

### EEG acquisition and processing

EEG data was acquired using a MR-conditional Brain Products system with a sampling rate of 5kHz and EEG acquisition software (Vision Recorder, B Brain Products, GmbH, Gilching, Germany) synchronized to the scanner clock and fMRI volume trigger (SyncBox, Brain Products, GmbH, Gilching, Germany). For all infants except one, data were acquired using a 32-channel cap adapted to newborns’ head size (Easycap GmbH, Wörthsee, DE). For one infant, we used a 25-channel cap due to the smaller head size and considered the correspondence between the common channels with the 32-channel cap.

MR gradient artefacts were removed from data using a sliding average template subtraction approach (Allen et al., 2000) in BrainVision Analyzer software (Brain Products, GmbH, Gilching, Germany). Before this step, recordings were visually pre-inspected to mark prolonged motion-artifacts periods and exclude these periods from template extraction. EEG data were low pass filtered at 40 Hz and down sampled to 250 Hz. As seen in previous datasets, cardiobalistic artifacts were not present in the majority of neonates (Arichi et al., 2017), except for two infants. For these infants, cardiobalistic artifact correction was performed using the same sliding average template subtraction approach, based on the information from the ECG channel recordings (Allen et al., 2000).

Data were then further preprocessed in MATLAB using EEGlab (Delorme & Makeig, 2004). Recordings were then filtered (0.5-15 Hz) and cleaned for motion artifacts using the APICE preprocessing pipeline (Flo et al., 2022). Briefly, this involved the identification and interpolation of persistently noisy channels, transient artifacts, as well as removal of motion-related contamination. We further applied independent component analysis to remove pulse artefacts, after which all recordings were visually inspected to eliminate remaining undetected artifacted periods. The data were re-referenced to the average reference, segmented into trials (i.e. epochs) time-locked to the onset of the first sound in the sequence (−100 to 3300 ms), and DC-detrended. Each trial activity was baseline corrected using the 100 ms window before stimulus onset, and trials were averaged for each sequence type to obtain auditory evoked responses.

#### Response habituation

Average evoked responses over frontocentral channels were used to extract the main response to the sound sequences. We first analyzed responses to the first three sounds (S1–S3) within sequences. Because these sounds were identical across all trials, standard, deviant, and omission trials were merged. For each infant, the amplitude rise between the N1 and P2 components of the auditory evoked potentials was quantified for each of the three sounds. To assess habituation following sound repetition, we then compared the amplitude difference between the three sounds with pairwise one-tailed t-tests and corrected for multiple comparisons using the false discovery rate (FDR).

#### Response to violation of regularity

To compare the response to deviant vs standard sounds, we focused on the [0-900] ms window after the onset of the last sound in these sequences. To compare the response to the unexpected omission with expected omission of sounds, we focused on [0-900] ms window post omission. This corresponded to unexpected omission of sound occurring in the 4^th^ place of omission sequences compared to expected omission occurring in the 5^th^ place of standard sequences. Since standard sequences are over-represented in an oddball paradigm, we selected only a subset for further comparisons with deviant/omission sequences. These corresponded to trials matching for occurrence position within blocks with respect to deviant/omission sequences (Adibpour et al., 2018). To identify the time-window where the two responses could differ, we performed non-parametric paired t-tests. Family-wise error correction was computed using a cluster-based permutation test (cluster-forming at alpha of 0.05, two-tailed t-tests with 5,000 permutations).

### MRI acquisition and processing

Data were acquired with a Philips Achieva 3T MRI scanner (Best, NL) and a 32-channel adult head coil, using an echo-planar imaging (EPI) sequence. A total of 540 volumes were acquired with 30 slices, TR=2001ms, TE = 37ms, FA = 90°, and resolution = 2.08×2.08×2.9mm, with a slice gap = 0.75 mm. Additionally, high resolution, 3D MPRAGE (Magnetization-prepared Rapid Gradient Echo) T1-weighted images with 0.8 mm isotropic resolution, TR/TI/TE = 11/1400/4.6 ms, SENSE factor 1.2 RL, were acquired for each infant to enable image registration to a template.

Image preprocessing was performed in FSL (www.fmrib.ox.ac.uk/fsl) (Woolrich et al. 2009). We retained only part of the functional scans without excessive and prolonged head motion (Dall’Orso et al., 2021) by computing the root mean square intensity difference between consecutive scan volumes to highlight major motion artifacts (Power et al., 2012) and then trimming the data to preserve the longest interval before or after intervals where more than 10 consecutive volumes were corrupted by motion artifact (corresponding to > 20 s of continuous motion). Three infants’ data were rejected at this step as it was only possible to retain less than 100 volumes. After trimming the fMRI data, the preprocessing included high-pass filtering (cut-off frequency = 0.01 Hz), MCFLIRT rigid body motion correction, slice timing correction and BET brain extraction, as well as spatial smoothing (Gaussian filter of FWHM 5 mm). Further denoising was performed to account for residual artifacts using independent component analysis whereby components manually labelled as artifact due to their temporal and spatial characteristics were regressed out prior to further analysis.

Voxel-wise General Linear Model (GLM) analysis was performed using FEAT (Smith et al, 2004). Each sound was modelled as an event, and deviant and omission events were orthogonalized with respect to repeated standard sounds to capture variance unique to these events. The study design was then convolved with a neonatal optimized basis function set (Arichi et al., 2012) to model task-related BOLD activity changes and define subject-level activation maps. To account for residual head motion–related artifacts, motion outliers were included in the GLM as regressors of no interest. z-scored F-statistical maps from each subject were first aligned to their own high-resolution anatomical image using rigid body registration and then warped to a T2-weighted 40-week infant template (Schuh et al., 2018) using ANTs (Avants et al., 2009). Group average effects were then identified using threshold-free cluster enhancement nonparametric testing, using one-sample sign-flipping permutations using randomise v2.1 (Smith & Nichols; 2009).

Although group-level activations to sound events are able to spatially localize the networks involved in processing, ongoing maturation of neurovascular coupling in the early developmental period means that Hemodynamic Response Function (HRF) shape features show significant variability across infants (Arichi et al., 2012; Moore et al., 2025). To account for this variability in the temporal features of the response, we performed exploratory epoch-based analyses on fMRI time-series extracted from each Region of Interest (ROI) identified from the group-level GLM analyses. We first z-scored the time series per subject and temporally upsampled the data with a factor of 20, using linear interpolation (Mazancieux et al., 2023). Time-series were then epoched around each block onset (time window: −2 to 40 seconds). To compare the BOLD signal across conditions, the baseline signal from the time window of −2 to 0 seconds was subtracted from each epoch. We then averaged the epochs per condition and participant. To assess the direction of change in BOLD activity for deviant/omitted sound events, we took the average amplitude of signal at the peak ±2 seconds and subtracted it from the BOLD activity in the standard blocks. To examine whether EEG responses to violations of regularity are associated with changes in fMRI BOLD activity in neonates, we used Spearman correlation to relate the interindividual variability in the EEG mismatch (i.e. average amplitude difference between deviant and standard sounds within identified time-windows of interest) to the peak amplitude of BOLD response for deviant versus standard sounds, after controlling for PMA at scan.

## Results

For EEG, infants (n=22) contributed a median of 84 trials (range: [20-105]). Within the group, 20/20/19 neonates contributed ≥5 trials per condition and had identifiable auditory evoked responses. In these infants, a total of 15 [6–23], 13 [6–17], and 13 [5–18] trials for standard/deviant/omission sequences were retained for the respective analyses.

### EEG responses to sound sequences

EEG auditory evoked potentials showed a distinct response could be identified following the presentation of each sound every 600 ms within a sequence. The main response components appeared first around 95 ms post-stimulus, with a negative polarity response (N1) followed by a positive polarity component (P2) appearing around 320 ms on the fronto-central channels. As described in the literature, there was a decrease in response amplitude from the first to the third sound (S1 → S3), visible on the time-course of the response, indicating habituation to repetition (Figure 2a). The peak-to-peak amplitude (N1 to P2 rise) for the first sound (S1) tended to be larger than the second sound (S1 vs S2: t (19) = 1.8, p= 0.060) and the third sound (S1 vs S3: t (19) = 2.4, p= 0.040). The second sound also tended to evoke a larger response than S3 (S2 vs S3: t (19) = 1.5, p= 0.079). There was no significant relationship between habituation degree and PMA across our infant subjects.

**Figure 2.**
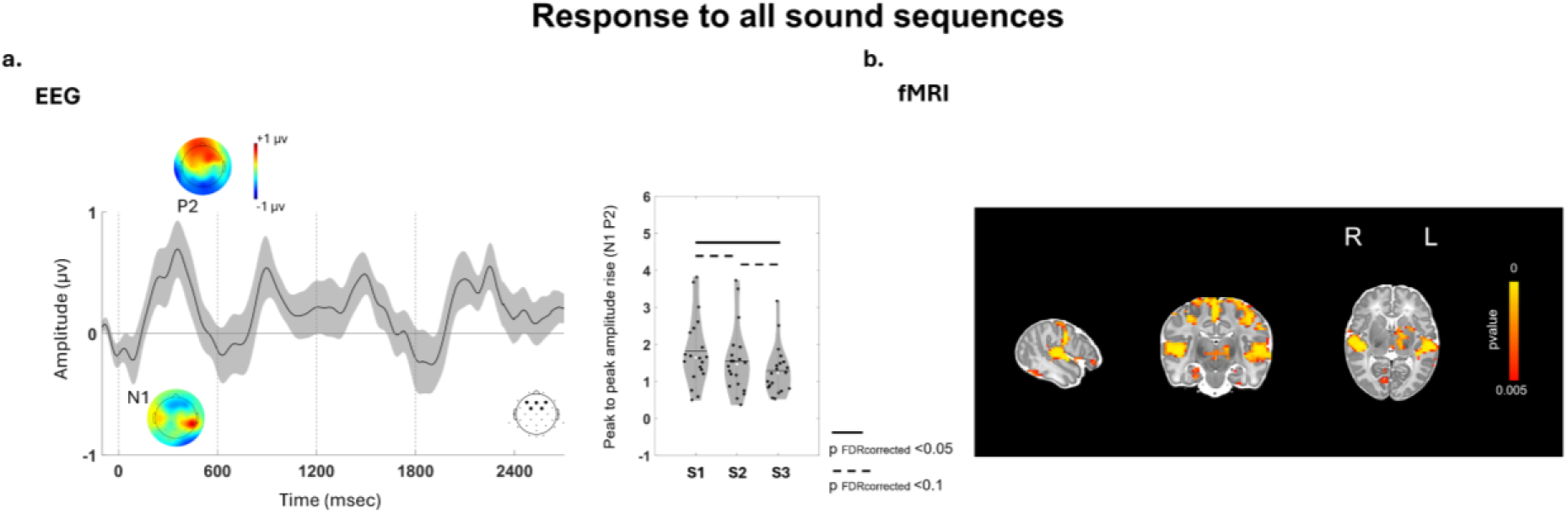
Main response to sound sequences. a) Average EEG activity over frontocentral channels across infants indicates sequential appearance of evoked potentials in response to the presentation of sound events. Dashed lines indicate the onset of each sound in the sequence. The topographical distribution of activity at the time of N1 and P2 components of the response are illustrated. Response amplitudes are compared between the first three sounds for each infant, indicating a general decrease in response from S1 to S3. b) fMRI group-level baseline activations to sound sequences indicate significant clusters of functional responses to sounds, which are projected to a 40-weeks infant template.

To characterise the specific response to violation of regularity, average EEG response on the frontocentral channels were compared between the deviant/omission and standard conditions. Cluster-based permutation analysis revealed a time-window with significant differences (p = 0.031) between the deviant vs. standard trials (Figure 3a). The difference corresponded a typical mismatch response similar to outside scanner recordings (Panzani et al., 2023), with an anterior positive pole (i.e. higher activations to deviant vs standard) and a reversal of polarity over the posterior/lateral regions. This delayed mismatch response was evident at 460-600 ms post stimulus period. There was no significant relationship between mismatch response and PMA across our infant subjects. No significant difference was observed for the comparison of unexpected omission vs. expected omission condition (Figure 3b).

**Figure 3.**
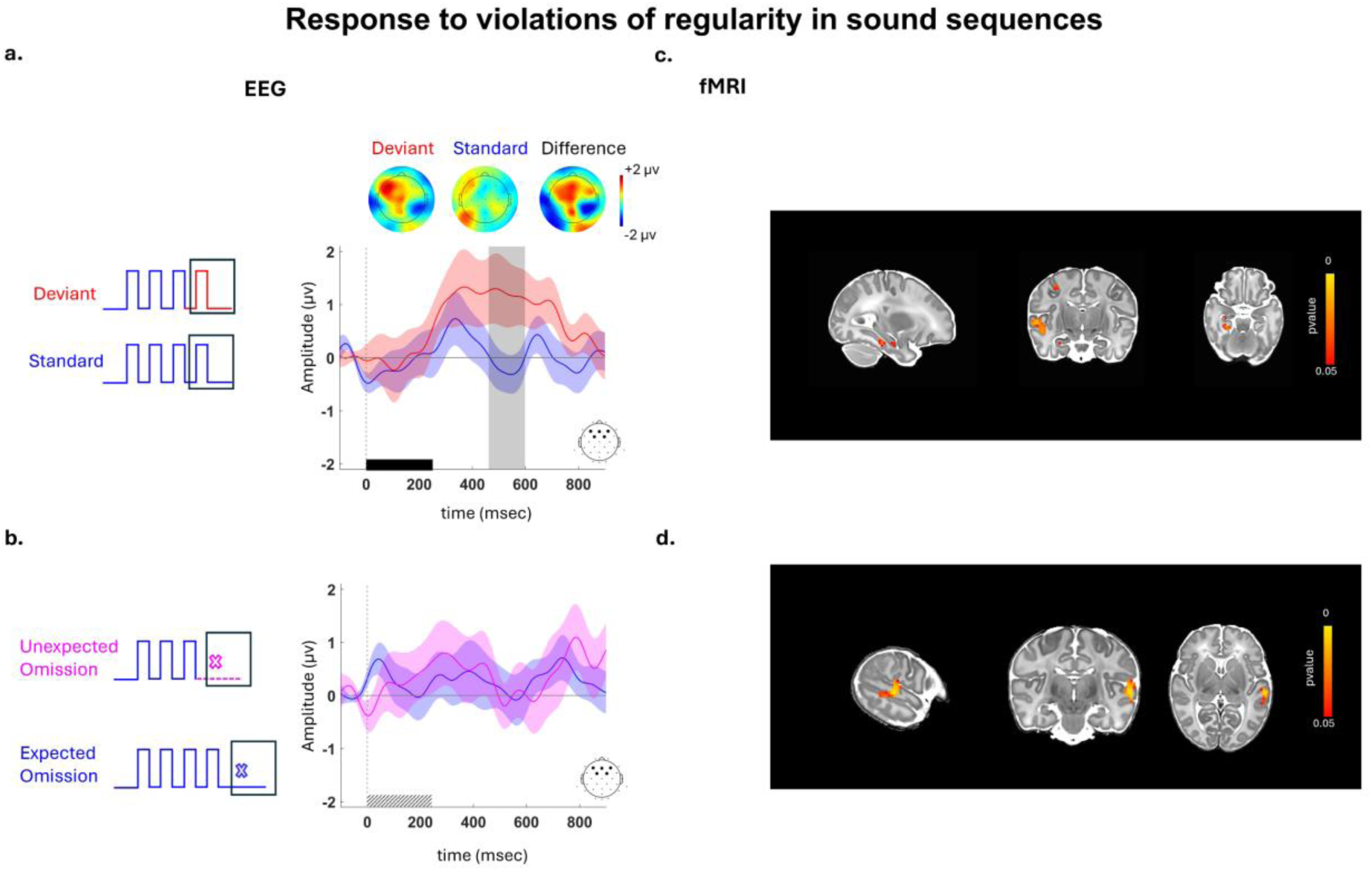
Response to violation of regularity in sound sequences. a) Average EEG activity over frontocentral channels across infants indicates significantly higher activations for deviant compared to standard sounds. b) The response to unexpected omission of sounds did not differ to that of expected omission of sounds at the end of the standard sequences. c) fMRI group-level activations to the occurrence of deviant sounds localized significant clusters of BOLD functional activity in the right superior and ventromedial parts of the temporal lobe, as well as precentral gyrus. d) fMRI group-level activations to the occurrence of unexpected omission localized significant clusters of BOLD functional activity in the left superior temporal gyrus.

For fMRI analyses, infants (n=22) contributed a median of 408 [151-540] fMRI scan volumes consisting of 10 [3-18] stimulation blocks. Out of these, 20/18/16 neonates contributed ≥2 blocks for each of the main experimental conditions (standard/deviant/omission) respectively and were retained for respective analyses.

### fMRI activations for sound sequence processing

GLM analyses identified significant clusters of BOLD functional activity in response to sound sequences, in a distributed network, encompassing the bilateral temporal, sensori-motor, and inferior frontal cortices, as well as subcortical regions within the left basal ganglia (including the pallidum and putamen), thalamus, hippocampus and parahippocampal areas (Figure 2b).

fMRI further localised significant clusters of activity in response to occurrences of deviant sounds in the right superior temporal gyrus and ventromedial parts of the temporal lobe, including the hippocampal and parahippocampal areas, as well as precentral gyrus (Figure 3c). Peak amplitude of the BOLD response within the identified regions were inspected to identify the direction of BOLD signal change for deviance as compared to standard sounds. All regions indicated a larger BOLD response peak for deviant compared to standard sounds (SI Figure 1a). Significant clusters of BOLD functional activity in response to occurrences of omitted sounds were localised in the left superior temporal gyrus (Figure 3d).

Interindividual variability in EEG mismatch responses was only marginally associated with peak BOLD activation to deviant sounds in the parahippocampal ROI after controlling for PMA at scan (SI Figure 2; r = 0.45, p = 0.053; p>0.7 for all other ROIs), but this relation did not survive multiple comparison.

## Discussion

Using simultaneous EEG-fMRI, we probed neonatal responses to sound sequences with regular and irregular structures, revealing both the time-course of the evoked activity and the spatial organization of the underlying networks. Our results suggest that neonatal brain detects violations of local, but not global, sequence structure relying on a distributed network spanning temporo–precentral areas and with the involvement of medial temporal subcortical structures.

### Adaptation to regularity and mismatch responses to violations of regularity

EEG analyses indicated reproducible peak latencies and topographies which were largely consistent with previous neonatal EEG studies examining the response to speech syllables (Dehaene-Lambertz and Pena, 2001; Mahmoudzadeh et al., 2017; Panzani et al., 2023), with the most distinguishable peak of the response appearing at approximately 320 ms. Similar to these studies, we saw a reduction in response amplitude with sound repetition within a sequence, suggesting that neonatal brain tracks the temporal regularity in repetition of consecutive sounds. A mismatch response was also observed following a violation of regularity in which the sound in the last position of the sequence distinctly differed from the preceding sounds. This response exhibited latencies and a topographical distribution comparable to those reported previously, in keeping with the detection of simple violations of regularity by sleeping neonates. Yet, no mismatch response was observed to the omission of sounds in the sequence.

Regularities and their violations can be represented and processed at multiple levels in the mature brain. Sequences can be encoded based on the frequency of individual items, the transitional probabilities between successive items, or more abstract rules about sequence structure such as the number and order of elements they contain (Dehaene et al., 2015). The observed reduction in responses with sound repetition is consistent with frequency-based encoding, often described as sensory adaptation (Ulanovsky et al., 2003). This adaptation may arise from neuronal refractoriness or from the sharpening of neural representations with repetition, such that fewer neurons respond to frequently occurring sounds.

Similarly, the neonatal mismatch response observed in our study may reflect at the most basic level, sensitivity to the low occurrence frequency of deviant sounds (frequency-based encoding), or sensitivity to the rare transition between the third sound and a deviant sound in the fourth position, consistent with neonates being capable of encoding transitional probabilities—often referred to as statistical learning (Saffran et al., 1996). Although the present paradigm does not allow these possibilities to be disentangled, previous work demonstrates that neonates are capable of encoding transitional probabilities in auditory sequences when frequency-based encoding is not sufficient to learn a structure (e.g., XYXYX structures; Panzani et al., 2023). Together, these findings suggest that probabilistic encoding of regularities (for individual items or the transition between items) may underly early auditory sequence processing.

By contrast, our results suggest that violations involving omissions in sequences may not be detectable by neonates using the same encoding strategies. In the more frequently presented standard sequences, the final sound is followed by silence, resulting in the transition from the last sound to silence occurring relatively frequently. As a result, omissions may not be detectable based on item frequency or transitional probabilities alone, but rather a representation of the global sequence structure, whereby the brain actively forms a prediction (i.e. expectation) (Summerfield & de Lange, 2014) about the number of items and presence of the last sound in the sequence. In line with this, prior work with adults have shown the presence of an early prediction response and a later mismatch response when an expected sound was omitted in a sequence (Wacongne et al., 2011; Heilbron, & Chait., 2018), suggesting that regularities induce precise temporal expectations based on sequence structure and when violated, mismatch responses could arise (Tal et al., 2017; Zalta et al., 2024).

The absence of mismatch response to sound omissions suggests that neonatal auditory encoding does not extend to global sequence structure, which in our paradigm would require tracking the number of embedded sounds (in this case 4). One possibility is that this reflects the immaturity of early brain networks, which could limit the capacity to encode higher-order global structure. In line with this interpretation, early numerosity perception in infancy appears to be very limited, in particular for more than 3 items (Coubart et al., 2013), potentially constraining the ability for a neonate to discriminate between sequences with three versus four items. Alternatively, the absence of an omission response could be related to state-dependent effect on neural processing as our neonates were studied in sleep. Although both our data and prior work show that the sleeping neonate brain can perform automatic comparisons between consecutive stimuli and detect simple and complex local deviances (Panzani et al., 2023), sleep may restrict the engagement of neural mechanisms required for processing higher-level sequences. In support of this view, a previous study has shown that sleep-stage differences in neonates modulate encoding of certain global-level properties in sound sequences (Moser et al., 2020). We could not assess the influence of sleep states on omission detection in our neonatal cohort, as recording time and the number of artifact-free trials per sleep stage were limited. This remains a direction for future investigation.

### Early networks underlying sound sequence processing within and beyond auditory cortex

fMRI analyses revealed that encoding sound sequences engaged a distributed network, encompassing subcortical structures as well as early and higher-order cortices beyond auditory areas. This network included the basal ganglia, thalamus, hippocampus and parahippocampal areas, as well as the superior and middle temporal gyri, the precentral gyri and the supplementary motor areas. These network level activations largely overlap with those reported during sound sequence learning in adults (Bekinschtein et al., 2009; Al Roumi et al., 2023). As the sequences in our paradigm inherently had a temporal regularity in their structure, their encoding likely also required processing of rhythmicity of stimuli. Consistent with this, the observed networks shared marked similarities with the cortico-subcortical networks described as being involved in musical rhythmicity perception in adults (Kasdan et al., 2022), particularly within cortical auditory and sensori-motor regions and subcortical nuclei, including the putamen and pallidum. Of relevance, recent infant fMRI studies have also found that sensitivity to structured and rhythmic auditory stimuli such as music is present very early in life. As in our study, activation is seen in infants up to 12 weeks of age within both primary and non-primary auditory regions of the temporal cortex, as well as in the hippocampi and amygdalae when listening to music (Perani et al., 2010; Kosakowski et al., 2023). Involvement of subcortical structures in early music processing has further been seen in preterm-born neonates, wherein early exposure to music was seen to alter functional connectivity between the thalamus, putamen and auditory cortex (Lordier et al., 2019). Moreover, the involvement of the supplementary motor area in early cross-modal learning (motor and auditory) has also been previously reported in neonates (Dall’Orso et al., 2021). Together, these findings point to an early emergence of functional pre-specialization of the infant brain for learning auditory regularities in the environment.

In our neonatal cohort, BOLD activity to deviant sounds were localized to areas beyond the primary auditory areas, highlighting a set of regions within superior and ventromedial parts of the temporal lobe, the latter encompassing the hippocampus and parahippocampus, as well as the precentral gyrus. This finding is in keeping with the hippocampus’ key role and early functional involvement in statistical learning (for detecting structures within visual arrays), as has been shown in awake infants at three months of age (Ellis et al., 2022). Our findings further point to early involvement of the hippocampus in encoding violations of regularity, despite its well-known protracted developmental trajectory (Uematsu et al., 2012).

Our results also complement the previous fNIRS studies showing responses to detection of regularity and deviances over temporal and frontal regions (Nallet et al.,2023; Mahmoudzadeh et al., 2013; Cabrera & Gervain, 2020). The whole brain coverage afforded by fMRI additionally enabled identification of involvement of the precentral gyrus and deeper nuclei such as hippocampus, which are potentially missed in fNIRS recordings given the restricted sensitivity to deep structures and limited sensor coverage. Note that recent fNIRS sensor arrays extending to sensori-motor regions, have enabled the observation of activation over these areas during detection of auditory regularities (Rajabi Mashhadi et al., 2025).

BOLD responses to the omission of sounds demonstrated activity over the left superior-temporal gyrus. A response to omission of a motor or somatosensory stimulus has been previously reported in neonates using fMRI and fNIRS (Dall’Orso et al., 2021; Dumont et al. 2017). However, since we did not observe a mismatch response in the EEG recordings, it is possible that the BOLD changes for omission may have captured the cumulative effect of response to consecutive sounds within blocks, with weaker positive BOLD to fewer sound events within omission blocks rather than a genuine response to detection of unexpected omission.

We observed only a weak relationship between individual mismatch responses and BOLD signal changes for deviant compared to standard sound sequences, with a trend indicating larger mismatch related to larger BOLD in parahippocampal areas. This may arise from the complexity of the maturational processes at this stage, as both cortical circuitry and the cerebral vasculature are still undergoing substantial maturation (Harris et al., 2011), with recent evidence indicating pronounced variations in BOLD response across cortical depth in neonates (Willers-Moore et al., 2025). Characterising the relationship between electrophysiological mismatch responses and fMRI BOLD activity in early infancy may therefore benefit from depth-resolved approaches to better delineate activity within laminar-specific circuitry and examine their relation to hierarchical processing of sensory regularity violations.

In sum, this study demonstrates the feasibility of investigating the temporal and spatial extent of early sensory activity in a few days’ old neonates, highlighting the precursor functional networks for processing auditory regularities and their violations. These networks, engaging cortical and subcortical regions, may provide the neural scaffolding that supports the early learning mechanisms and later emergence of complex cognitive abilities, including language learning.

## Acknowledgments

We thank all infants and families who agreed to participate in this work, and the staff of St. Thomas’ Hospital London. P.A. was supported by the UK Government Horizon Europe funding guarantee scheme of MSCA [EP/X021947/1]. T.A. was supported by an MRC Clinician Scientist Fellowship [MR/P008712/1] and a Transition Support Award [MR/V036874/1]. P.A and T.A. were additionally supported by an MRC Senior Clinical Fellowship [MR/Y009665/1]. The work was also supported by a project grant awarded by Action Medical Research [GN2728]. J.O.M. was funded by a Sir Henry Dale Fellowship jointly by the Wellcome Trust and the Royal Society [206675/Z/17/Z]. T.A. and J.O.M. received support from the Medical Research Council Centre for Neurodevelopmental Disorders, King’s College London [MR/N026063/1].

## Supplementary Information

**Supplementary Figure 1.**
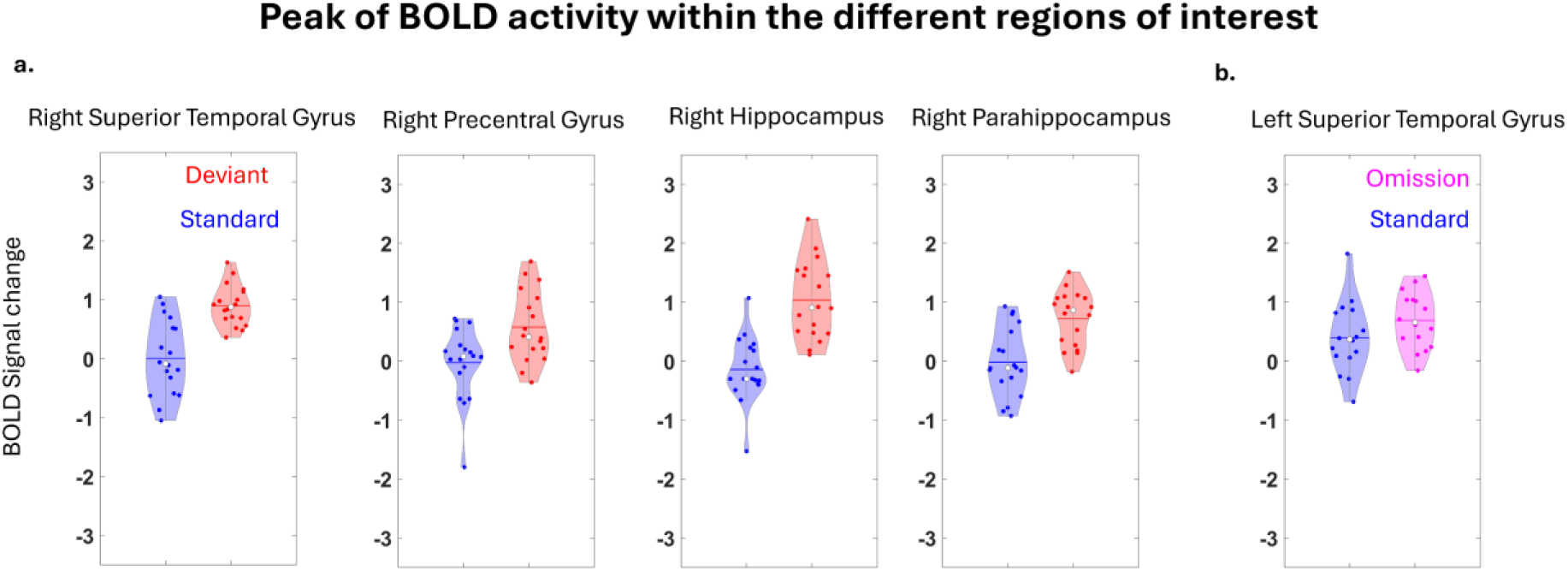
Peak amplitudes of the BOLD response within regions identified in group-level fMRI analyses for the occurrence of a) deviant sound and b) sound omission.

**Supplementary Figure 2.**
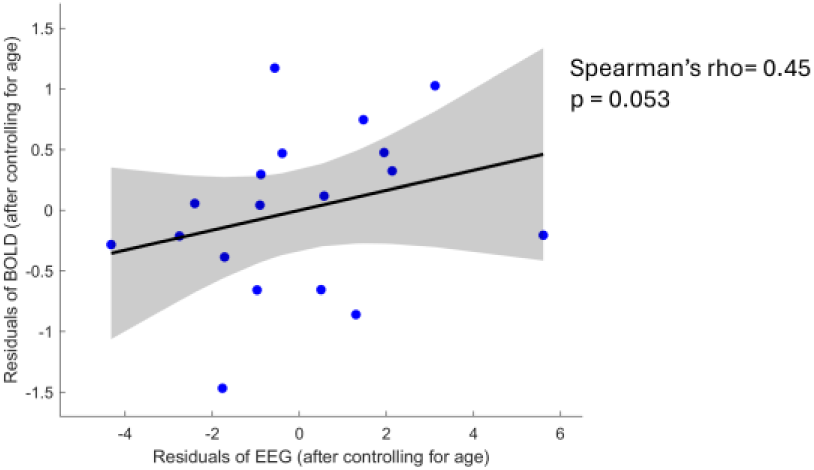
Interindividual variability in the peak amplitudes of the BOLD response to deviant versus standard sounds were related to the amplitude of EEG mismatch response after regressing out PMA at scan.

